# Fusion of DARPin to aldolase enables visualization of small protein by cryoEM

**DOI:** 10.1101/455063

**Authors:** Qing Yao, Sara J. Weaver, Jee-Young Mock, Grant J. Jensen

## Abstract

In recent years, solving protein structures by single particle cryogenic electron microscopy (cryoEM) has become a crucial tool in structural biology. While exciting progress is being made towards the visualization of smaller and smaller macromolecules, the median protein size in both eukaryotes and bacteria is still beyond the reach of single particle cryoEM. To overcome this problem, we implemented a platform strategy in which a small protein target was rigidly attached to a large, symmetric base via a selectable adapter. Seven designs were tested. In the best construct, a designed ankyrin repeat protein (DARPin) was rigidly fused to tetrameric rabbit muscle aldolase through a helical linker. The DARPin retained its ability to bind its target, the 27 kDa green fluorescent protein (GFP). We solved the structure of this complex to 3.0 Å resolution overall, with 5 to 8 Å resolution in the GFP region. As flexibility in the DARPin limited the overall resolution of the target, we describe strategies to rigidify this element.

**Author summary:** Single particle cryogenic electron microscopy (cryoEM) is a technique that uses images of purified proteins to determine their atomic structure. Unfortunately, the majority of proteins in the human and bacterial proteomes are too small to be analyzed by cryoEM. Over the years, several groups have suggested the use of a platform to increase the size of small protein targets. The platform is composed of a large protein base and a selectable adapter that binds the target protein. Here we report a platform based on tetrameric rabbit muscle aldolase that is fused to a Designed Ankyrin Repeat Protein (DARPin). Phage display libraries can be used to generate DARPins against target proteins. The residues mutated in a phage display library to generate a DARPin against a new target do not overlap with the DARPin-base fusion in the platform, thus changing the DARPin identity will not disrupt the platform design. The DARPin adapter used here is capable of binding Green Fluorescent Protein (GFP). We report the structure of GFP to 5 to 8 Å local resolution by single particle cryoEM. Our analysis demonstrates that flexibility in the DARPin-aldolase platform prevents us from achieving higher resolution in the GFP region. We suggest changes to the DARPin design to rigidify the DARPin-aldolase platform. This work expands on current platforms and paves a generally applicable way toward structure determination of small proteins by cryoEM.

## Introduction

Single particle cryoEM can reveal the structures of large macromolecular complexes to near atomic resolution. To solve a protein structure by single particle cryoEM, purified proteins are rapidly frozen in a thin layer of vitreous ice. A transmission electron microscope is used to collect projection images of the protein. Individual proteins are identified in the ice and their orientations are computationally determined. The projection images are then combined to calculate a 3D reconstruction of the protein.

A fundamental challenge in single particle cryoEM is that small proteins do not produce enough contrast in noisy projection images to precisely determine their orientation. Richard Henderson estimated that with ideal images, a 3 Å structure could be reconstructed for a 40 kDa protein (1). Unfortunately, real electron micrographs are imperfect so this theoretical minimum of macromolecular size has never been reached. The smallest protein to be solved to near atomic resolution so far by cryoEM is hemoglobin (64 kDa) (2), but the median protein lengths in both bacteria (27 kDa) and eukaryotes (36 kDa) are about two times smaller (3). Consequently, many proteins in biology are beyond the reach of high-resolution structure determination by single particle cryoEM.

Over the years, several strategies to overcome the size limit problem in single particle cryoEM have been suggested. Two major themes have emerged to increase the target mass and improve its orientation determination. First, the target can be decorated with antibody fragments (6) (7). Second, the target can be rigidly attached to a large platform protein. The platform is typically composed of a base protein and an adapter. The purpose of the base protein is to increase the molecular weight, which facilitates accurate particle picking and precise particle orientation determination. The adapter can be customized (a covalent fusion between the target and the base) or general (a selectable adapter that facilitates non-covalent binding of the target to the platform base). Covalent approaches have utilized direct fusions between the target protein and the base either via a flexible linker adapter (4) or a helical junction adapter (5) (8). For the platform to be successful, the adapter must be rigidly attached to the base. The flexible linker adapter was therefore insufficient to determine the structure of the target (4), but the use of a helix-forming peptide linker (8)(9) or direct concatenation of two helices (5)(10) has shown promise. Most recently, Liu et al. demonstrated that a rigid, continuous α-helix could be formed by linking the terminal α-helices of a designed ankyrin repeat protein (DARPin) and a nanocage subunit through a helix-forming peptide linker (9) (8). Notably, Liu et al were able to show the structure of the 17 kDa DARPin to 3.5 to 5 Å local resolution (8). Unfortunately, these strategies are limited to target proteins with a terminal α-helix, and their implementation requires that the length of the helical junction adapter must be customized for each new target. Utilizing a non-covalent platform strategy with a selectable adapter (like an antibody or a DARPin) has the potential to be generally applicable, regardless of the structure of the target, since the selectable adapter could be raised against any target using phage display, while the invariant nature of the adapter framework region would allow the one-time optimization of a rigid attachment point between it and the base. Along these lines, Liu et al. suggested that their DARPin-nanocage could display a small protein for structure determination by cryoEM (8), but so far no group has demonstrated this.

Here we report the outcomes of a variety of new designs and report the structure of the first small protein visualized through a base/selectable-adapter platform approach.

## Results

### Platform strategy and design

The goal of our study was to design a generally applicable platform to solve small protein structures by single particle cryoEM. We explored several candidate base proteins and selectable adapters (Fig S1). We favored bases that were easy to purify and that had already been solved to high-resolution by single particle cryoEM. We reasoned that oligomeric and symmetric (as a globular protein, or as a helical tube) bases would be best.

As selectable adapters, we first considered antibody fragments (Fabs and scFvs). Fabs have a flexible elbow connecting two immunoglobulin regions, whereas scFvs are made up of one immunoglobulin region. The Fab elbow could introduce flexibility, so we preferred the smaller, more compact scFv. However, because the beta sandwich immunoglobulin fold of a scFv could be difficult to rigidly fuse to the surface of a platform base, we identified Staphylococcus Protein A (PrA) as a linker that could bind the invariant region of a scFv (11). As PrA is a three-helix bundle, we reasoned that it could be rigidly attached to a base via a helical linker. Thus in one of our designs, the C-terminal helix of PrA was fused to the N-terminal helix of the base protein. Since PrA is capable of binding the invariant scFv framework, the base-PrA:scFv interfaces would not need to be redesigned for each new target. Unfortunately, in our biochemical experiments, we observed that the PrA:scFv interaction did not remain stable through a gel filtration column, indicating that the binding affinity was not strong enough for our purposes. Further mutagenesis to the PrA:scFv interface may strengthen the interaction. Regardless, a fundamental concern with this design is that two non-covalent binding interactions are required (PrA:scFv, and scFv:target), which could lead to occupancy issues. As a result, we moved to DARPins as our selectable adapter (Fig 1B).

**Fig 1.**
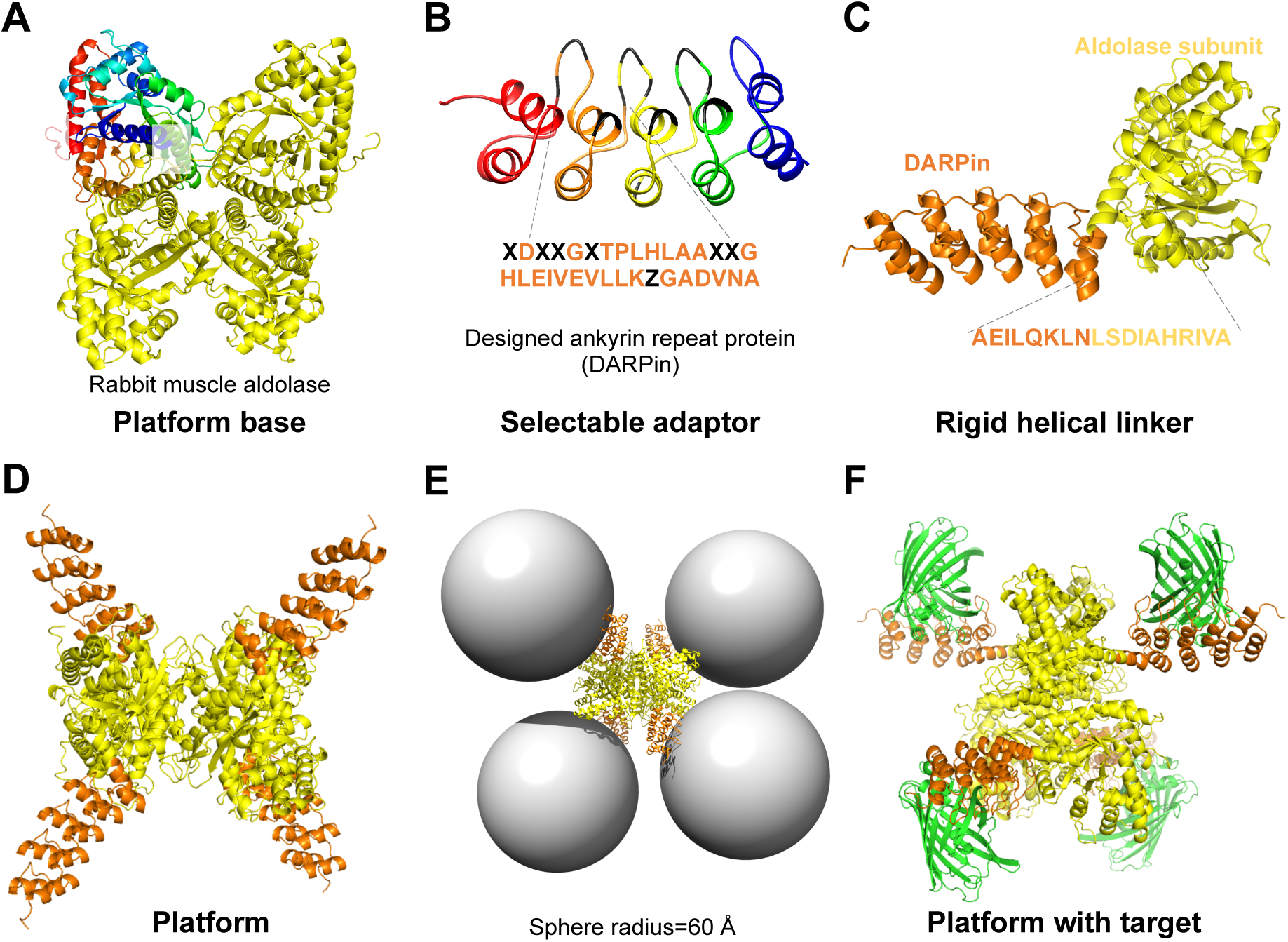
The Design of the platform. (A) The platform base was homotetrameric rabbit muscle aldolase (PDB ID code 5VY5). One subunit was depicted with rainbow coloring and N and C labels to indicate the orientation of the monomer chain. The other three identical subunits are shown in yellow. Aldolase has D2 symmetry. (B) The selectable adapter was a Designed Ankyrin Repeat Protein (DARPin) (PDB ID Code 5MA6). A DARPin is made up of a series of ankyrin repeat motifs (a beta turn followed by two antiparallel alpha helices). Here a DARPin against GFP was used. The general structure of the library this DARPin came from is an N-terminal cap (red) followed by a series of internal binding modules (here module 1 (orange), module 2 (yellow) and module 3 (green)) that are stabilized by a C-terminal cap (blue) (12). Shown below is a close-up view of the repetitive motif of DARPin with its amino acid sequence (orange). Using a phage display library, DARPins can be generated against a protein target. The selectable residues are depicted in black as X (any amino acid except cysteine or proline) or Z (amino acids asparagine, histidine or tyrosine) (12). (C) The final helix of the C-terminal cap of the DARPin (orange) was directly fused to the first alpha helix of aldolase (yellow) to form the platform subunit. (D) The D2 symmetry of the DARPin-aldolase fusion demonstrates ample space for target binding. (E) Spheres (radius=60 Å) were drawn in the position where each DARPin binds its target. A globular protein of up to 740 kDa could be accommodated on the DARPin-aldolase platform without steric clash. (F) The model of the DARPin-aldolase platform in complex with GFP (green) is shown.

In our designs, the final alpha helix of the DARPin C-terminal cap (C-cap) was directly fused to the first α-helix of the base (Fig 1C). All DARPin libraries use a C-cap to stabilize the protein, so we expect it will be straightforward to swap in any DARPin built on the same framework (Fig 1B). In the base-DARPin platform design, only one non-covalent interaction is required (between the DARPin and the target), which results in a more predictable and stable complex. We chose a DARPin that formed a stable complex with GFP as a first test case (12) and screened several base-DARPin candidates.

### Screening base candidates

We performed expression trials for five of our base-DARPin candidates (Fig S1). These bases included β-galactosidase (β-gal) (13), the vipA/vipB helical tube (14), the *E. coli* ribosome (15), TibC (16), and aldolase (17).

Because β-gal tetramerization requires the N- and C-termini of each subunit (18), an internal DARPin insertion was used, flanked by a helix-forming peptide (at the DARPin N-cap) and a flexible linker (at the DARPin C-cap) (9). Biochemically the ß-gal-DARPin platform formed a stable complex with GFP, but no cryoEM density was observed for the DARPin or GFP in our 3 Å reconstruction. This means that the helical linker was flexible relative to the β-gal base.

We therefore focused on bases with a terminal α-helix that could be rigidly fused to the DARPin. The vipA/vipB, ribosome L29, TibC, and aldolase proteins all had long terminal α-helices to facilitate direct fusion. In our experiments, the helical tube vipA-DARPin/vipB platform exhibited poor expression in *E. coli*, while the L29-PrA fusion did not integrate well into ΔL29 *E. coli* ribosomes (15) (Fig S1). The purified TibC-DARPin platform formed a stable complex with GFP, but the complex demonstrated aggregation and preferred orientation on plunge frozen grids. In contrast, the DARPin-aldolase platform was well-behaved.

In our DARPin-aldolase platform, the C-terminal α-helix of the DARPin was directly concatenated to the N-terminal α-helix of aldolase (Fig 1C) (S2 Fig A). The D2 symmetry of the DARPin-aldolase platform provided extensive space for the target and could potentially accommodate a globular protein of up to 740 kDa without steric clash (Fig 1E, 1F) (S1 Movie). The purified GFP:DARPin-aldolase complex was stable in a gel filtration column with an apparent 1:1 stoichiometry of DARPin-aldolase to the target (GFP) (S2 Fig B and C).

### CryoEM analysis of the GFP:DARPin-aldolase complex

To solve the structure of GFP bound to the DARPin-aldolase platform, we collected 1,681 micrographs on a Titan Krios (Fig S3). Because the thin ice forced a slight preferred orientation issue, an additional 1,180 micrographs were collected at 26° tilt (see methods) (19). High quality micrographs were selected after CTF correction (Fig S4) and particles were autopicked in Relion (Fig S3). After 2D classification in cryoSPARC, classes with strong secondary structure were selected for reconstruction. The GFP:DARPin-aldolase complex reconstruction yielded an overall resolution of 3 Å with C1 symmetry (Fig S5B). Further classification suggested too much conformational heterogeneity to apply D2 symmetry. The aldolase core and the helical linker were resolved to near atomic resolution (Fig 2B, 2C, 2D). The DARPin and GFP exhibited a local resolution of 4 to 8 Å, with discontinuous regions of higher resolution of 3.5 Å (Fig 2D) (S2 Movie). Although the resolution in the GFP and DARPin portion was not sufficient to build a model or assign sequence *de novo*, the static X-ray structures of GFP and the DARPin could be reliably docked into the map (Fig 2A).

**Fig 2.**
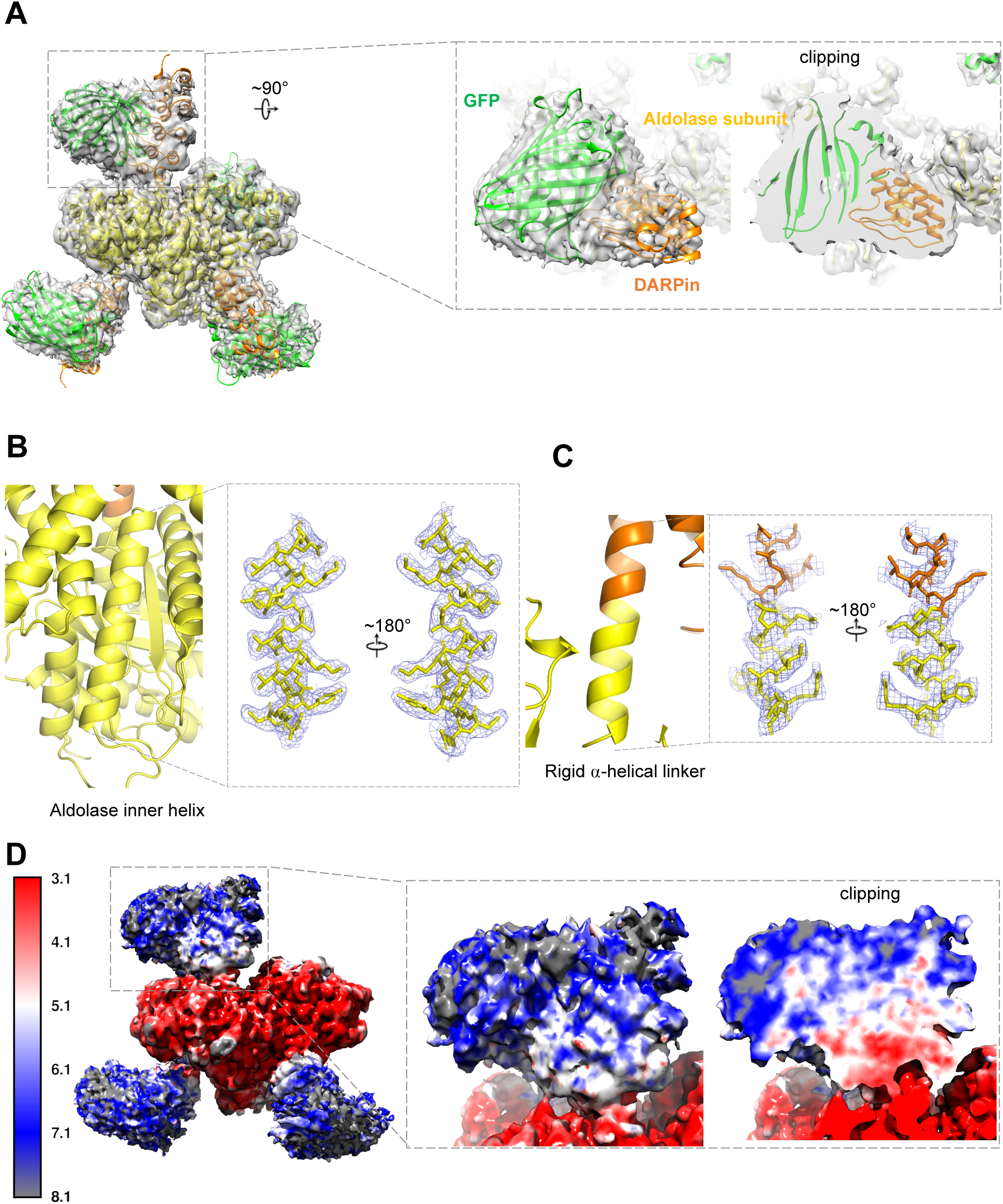
The cryoEM structure of the DARPin-aldolase platform in complex with the target GFP. (A) Surface of the 3 Å C1 reconstruction of DARPin-aldolase platform in complex with GFP. The crystal structures of GFP (green), the DARPin (orange) and aldolase (yellow) were docked into the cryoEM density. Shown on the left is the overall structure and on the right is the expanded view of GFP and the DARPin fit into the density of the best resolved subunit. The expanded view in the dashed line box is shown from the top (left) and halfway down the DARPin with clipping (right) to indicate the quality of the fit. (B) Ribbon diagram (left) and cryoEM density (right, blue mesh, zoned 1.8 Å within atoms) of an internal aldolase helix (residues Arg369 to Asp387). (C) Ribbon diagram (left) and cryoEM density (right, blue mesh, zoned 1.8 Å within atoms) of the helical linker (residues Ala176 to Ile191) between the DARPin (orange, residues Ala176 to Lys181) and aldolase (yellow, residues Leu182 to Ile191). (D) ResMap local resolution estimate of the final DARPin-aldolase platform in complex with GFP (left) and of the best subunit (right). The expanded view in the dashed line box is shown from the side (left) and halfway into the GFP:DARPin density with clipping (right).

### DARPin framework caused conformational heterogeneity

Because of the 5 to 8 Å local resolution range in the GFP portion of the map (Fig 2D), we suspected that part of the GFP:DARPin-aldolase complex was flexible. To better understand the conformational heterogeneity in the data, a mask was generated around a single DARPin/GFP unit and Relion particle symmetry expansion was used to consider each subunit individually (Fig 3A, Fig S3) (21). The symmetry expanded particles were subjected to 3D classification without alignment, a strategy in which the orientation parameters determined in the previous refinement are used to classify the particles into subsets. For this focused classification, a spherical mask that encompassed the aldolase surface was used to increase the signal. The resulting five classes showed reasonable GFP:DARPin conformations (Fig 3B), but subsequent refinements were still limited to 5 to 6 Å overall, which suggested that additional conformational heterogeneity remained within the subsets. The majority of the particles (54%) were classified into class 2 (yellow), which appeared to lack a DARPin (Fig 3B). Class 2 was subjected to an additional round of 3D classification where it revealed several reasonable but lower resolution GFP:DARPin conformations (Fig S6). To investigate the heterogeneity in the focused classes, we compared each class to class 4 (Fig 3C, 3D). In the different classes, the GFP:DARPin density shows a clear rocking around the Y axis (Fig 3C) and around the Z axis (Fig 3D) relative to the aldolase base. At this point, we wondered if any these displacements could be attributed to the aldolase subunit. We performed a similar focused classification experiment with a mask around the aldolase subunit and the helical linker, but no rotation or shift was observed in the resulting subsets (data not shown). Thus, we concluded that the displacement likely arose in the C-cap second helix that is fused into the helical linker, and other regions of the DARPin distal to the linker.

**Fig 3.**
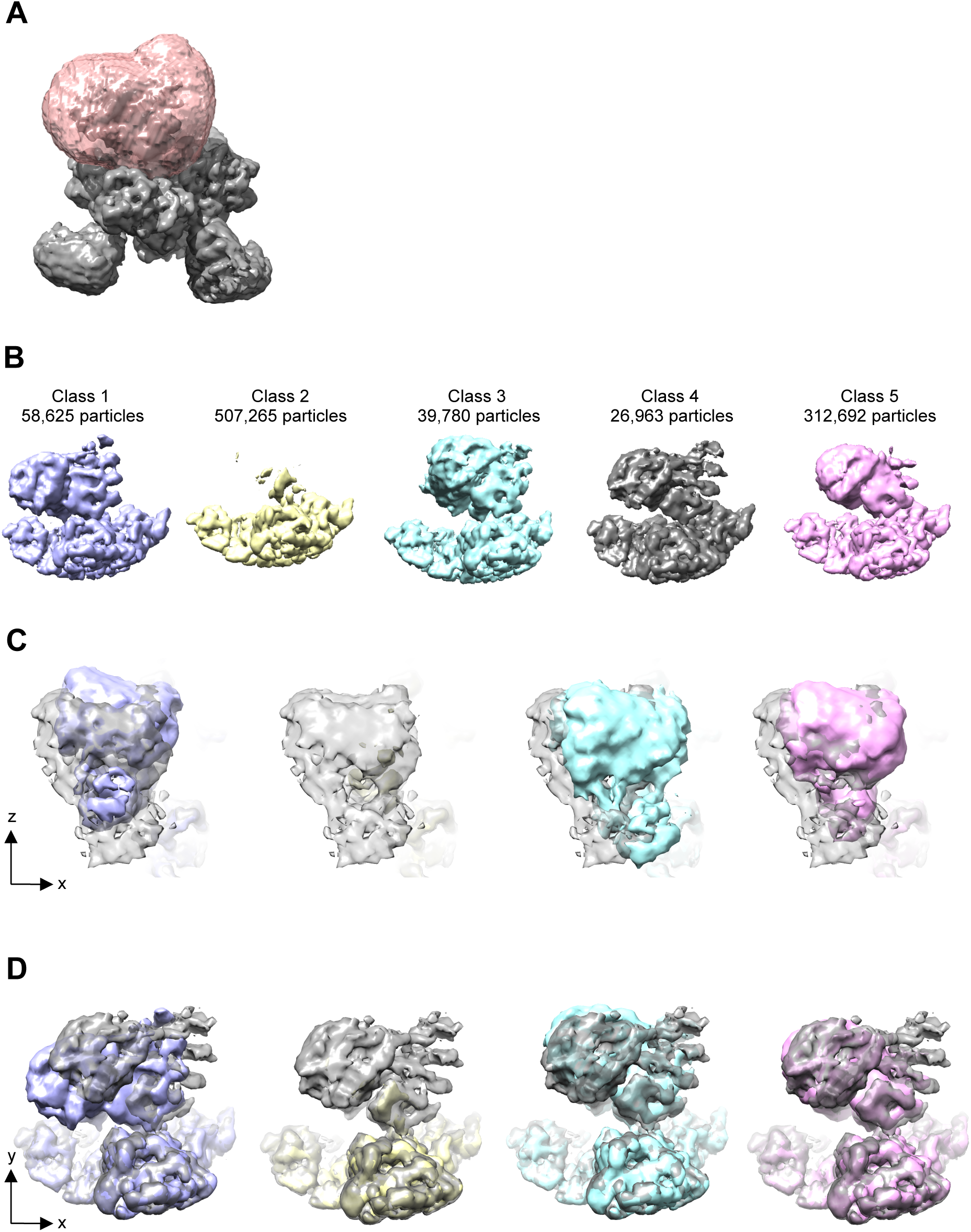
Symmetry expanded 3D classification of the GFP:DARPin region of the density. (A) CryoEM density of the D2 reconstruction of the DARPin-aldolase platform in complex with GFP (grey) is shown with a mask (pink) around one GFP:DARPin region. Symmetry expansion was applied to this particle set so that each GFP:DARPin region in each tetramer could be considered independently in 3D classification. (B) 3D classification without alignment of the symmetry expanded particles yielded five reasonable classes. The number of particles per class is indicated above each class. Each density was viewed at the same threshold in Chimera to facilitate direct comparisons. (C) The classes in Fig 3B were each compared with Class 4 (grey) to show the displacement between classes. The XZ plane is shown and the Y axis is perpendicular to the page. Class 4 was clearly displaced relative to the other classes. (D) The comparison from Fig 3B is now viewed looking down the Z axis. (E) The view from Fig 3D was adjusted to look down the helical linker. Residue Arg190, the C-cap helix 1, and the binding module 3 are labeled with arrows to orient the reader relative to Fig 1.

## Discussion

In this study, we designed and tested a variety of platforms capable of non-covalently binding a small target protein via a selectable adapter for structure determination by single particle cryoEM. In our best construct, we resolved our target protein (GFP) to 5 to 8 Å resolution.

Our DARPin-aldolase platform has several advantages over other strategies. It is simple to express and purify. Aldolase has D2 symmetry and allows attachment of four targets without steric clash. Aldolase can be reconstructed to 2.6 Å resolution with even a 200 keV microscope (17). Because DARPins can be readily generated against a wide range of small protein targets, the attachment of a DARPin to aldolase promises to be a generally applicable strategy. A recent study of the insulin degrading enzyme (IDE) bound to Fabs was able to isolate several IDE conformations using different Fabs (22). It stands to reason that different DARPins could also stabilize different conformations of the target. Because switching DARPins in the platform would be done by straight-forward DNA manipulations, our DARPin-aldolase platform has the potential to resolve a series of conformations of the target protein.

Our biochemistry experiments suggested that the purified GFP:DARPin-aldolase complex was very stable, and clear secondary structure was apparent in the 2D classes, yet heterogeneity remained. Because the aldolase base and the helical linker region were resolved to near atomic resolution (Fig 2B and 2C), the heterogeneity likely began in the DARPin C-cap. The DARPin against GFP used here was from a first generation DARPin library. The C-cap of the first generation DARPins was reported to be less stable than the other repeat modules (23). While the crystal structure contained a well-resolved C-cap, the heterogeneity observed here suggests that it is not yet sufficiently rigid to serve as an attachment point in a cryoEM platform (Fig 3). Recent DARPin phage display libraries contain DARPins with reduced surface entropy and a more stable C-cap sequence (23), however, and additional stabilizing surface interactions could be introduced in future designs (28) (29), or even a second attachment point of the DARPin to the base (at both N- and C-terminal caps of DARPin for instance). Together such improvements could allow the DARPin-aldolase platform to reveal the structures of many small proteins to near atomic resolution.

## Materials and Methods

### Computational design

Computational α-helix fusion was generated by manually docking the rabbit muscle aldolase structure (PDB code: 5VY5) and GFP/DARPin complex (PDB code: 5MA6). In order to rigidly join the aldolase and DARPin moiety together, we truncated the C-terminal flexible loop on DARPin and N-terminal flexible loop on aldolase, respectively, exposing the two terminal α-helices. The two terminal α-helices were manually concatenated and joined together to form an ideal α-helix using building α-helix tool in UCSF Chimera (30). The model was inspected for the orientation of DARPin relative to the aldolase, ensuring no steric clash and the providing enough space for target protein attachment. All structural design figures were generated using PyMOL1.8 (https://pymol.org).

### Cloning, protein expression, and purification of the recombinant DARPin-aldolase platform and GFP

The DARPin sequence was DARPin 3G86.32 (Fig S2A) (12). The cDNA expressing GFP and our DARPin-aldolase fusion were synthesized at IDT DNA company. The cDNA of GFP and DARPin-aldolase fusion were PCR-amplified and inserted into pACYCDuet and pET21b vector for recombinant expression in *E. coli*, producing no-tag GFP protein and C-terminal His-tag of DARPin-aldolase chimeric fusion. GFP and DARPin-aldolase were coexpressed in *E. coli* BL21(DE3) (Lucigen) using autoinduction medium with trace elements (Formedium) at 30 °C for overnight. Cells were harvested by centrifugation and the protein complex was then purified with Ni-NTA affinity chromatography (Qiagen), and Superdex 200 chromatography (GE healthcare). The purified GFP-DARPin-aldolase complex was concentrated to 2.5mg/ml in a buffer containing 25 mM Tris-HCl pH 8.0 and 150 mM NaCl.

### CryoEM sample preparation and data collection

Electron microscopy grids were prepared at Scripps Research Institute. Briefly, 3 µL sample of 2.5 mg/ml GFP-DARPin-aldolase complex was applied to a plasma cleaned Au UltraFoil Grid (300 Mesh, R2/2, Quantifoil) in a cold room (4°C, ≥95% relative humidity). The grid was manually blotted with a filter paper (Whatman No.1) for approximately 3 seconds before plunging into liquid ethane using a manual plunger (17). The grids were screened in Talos Arctica 200 kV with Falcon 3 (FEI) direct electron detector for ice thickness and sample distribution. Micrographs of GFP-DARPin-aldolase complex were collected on Titan Krios microscope (FEI) operating and 300 kV with energy filter (Gatan) and equipped with a K2 Summit direct electron detector (Gatan). For untilted data, Serial EM was used for automated EM image acquisition (31). After calculating an efficiency score from early refinements using cryoEF (19), additional data were collected at 26° using EPU software (FEI). A nominal magnification of 165,000x was used for data collection, corresponding to a pixel size of 0.865 Å at the specimen level, with the defocus ranging from -1.0 µm to -3.0 µm. Movies were recorded in superresolution mode, with a total dose of ~40 e-/ Å^2^, fractioned into 20 frames (0° tilt images) or 40 frames (26° tilt images) under the does rate of 8.4 electron per pixel per second.

### Image processing and structure analysis

Movies were decompressed and gain corrected with IMOD (32). Motion correction was performed using program MotionCor2 (33), and exposure filtered in accordance with relevant radiation damage curves (34). Micrographs with high CTF Figure of Merit scores and a maximum resolution great than 3.6 Å were selected for further processing. Particles were autopicked using 2D classes as references and extracted in RELION (35) and initial 2D classification was performed in cryoSPARC (36). High quality 2D classes were selected for further processing. The initial model was *de novo* generated and subsequent 3D refinement were performed using cryoSPARC. The UCSF PyEM package (*https://github.com/asarnow/pyem*) script was used to convert the cryoSPARC coordinates into Relion. Duplicate particles were removed and particles were analyzed by 3D refinement, Bayesian Particle Polishing and CTF Refinement in Relion. The data were binned to 1.5 Å/pixel, refined with D2 symmetry, and symmetry expanded. Symmetry expanded particles were used in 3D classification without alignment. All reconstructions were analyzed using USCF Chimera. The initial model was built rigidly docking individual protein structures into the EM map using Chimera. The model was then fit and adjusted manually in USCF Chimera and Coot (37). The figures were generated using UCSF Chimera, and local resolution and final Fourier shell correlation were calculated using ResMap (38) and cryoSPARC.

### Data deposition

Density map of GFP:DARPin-aldolase complex has been deposited in the Electron Microscopy Data Bank (EMDB) with access code: EMD-9277 and PDB 6MWQ.

## Acknowledgement

We thank Dr. Mark Herzik Jr. and Mengyu Wu at The Scripps Research Institute for help with sample preparation. We also thank Dr. Songye Chen, Dr. Andrey Malyutin, Dr. Rebecca Voorhees, and Dr. Bil Clemons at Caltech for technical assistance. We are also grateful to all members of the Jensen laboratory for discussion and technical assistance. This work was supported by funds from NIH NIGMS P50 082545. CryoEM work was performed in the Beckman Institute Resource Center for Transmission Electron Microscopy at Caltech.

## Supporting information

**S1 Fig. Attempted platform designs and outcomes**

(A) Models of the seven platform base proteins tested here. Different subunits of the platform were drawn by different colors. The fused DARPin (in green) and target GFP (in cyan) were shown for only one subunit for clarity. (B) Table summarizing the progress and problems related to our designs

**S2 Fig. Sequence of DARPin-aldolase fusion and the purification of GFP-DARPin-aldolase complex**

(A) The amino acid sequence of DARPin-aldolase fusion is colored with the DARPin sequence in brown and the aldolase sequence in green. The residues that were randomized in the phage display library are colored in red. The secondary structures are indicated on top of the sequence with α-helix in magenta tubes and β-strand in green arrows. The rigid helical linker is represented by a blue tube. (B) Gel filtration chromatography of the purified DARPin-aldolase platform in complex with GFP on Superdex 200 column. The black arrows mark the molecular weight calibration and void volume. Fractions 1 to 5 are labeled. (C) SDS-Page gel stained with Coomassie Blue of fractions 1 to 5 from the gel filtration chromatography. The bands representing the DARPin aldolase platform subunit and the GFP are labeled.

**S3 Fig. Major steps of the last cycle of cryoEM data processing**

**S4 Fig. Raw micrographs with CTF correction at 0**° **and 26**° **tilt**

(A)-(C) Motion-corrected, dose weighted micrograph of DARPin-aldolase platform in complex with GFP in vitreous ice is acquired at a nominal magnification of 165,000× (left) with the Fourier transformation (inset) and the CTFFind4 plot result (right). Micrographs were collected at 0° tilt in the first session (A), at 0° tilt in the second session (B) and at 26° tilt in the third session (C). Each micrograph has been low-pass filtered to 10 Å to enhance the contrast. The power spectrum of this micrographs is shown as an inset. CTF estimation fit (orange line) to experimental power spectrum (green line) and quality of fit (blue line) are plotted against spatial frequency (1/Å). Scale bar, 20 nm.

**S5 Fig. 2D classes and FSC from the DARPin-aldolase platform in complex with GFP refinement**

(A) Representative 2D classification results from Relion. (B) Relion Post Processing Fourier Shell Correlation (FSC) plot for the C1 refinement of the DARPin-aldolase platform in complex with GFP. A B factor of -75 Å^2^ was used to sharpen the map. FSC for the phase randomized masked map (red), the unmasked map (green), the masked map (blue) and the corrected map (black) are plotted. The FSC=0.143 cutoff is annotated with a black horizontal line.

**S6 Fig. 3D classification without alignment of the symmetry expanded class 2.**

Class 2 (yellow, top row) from the 3D classification discussed in Fig 3 contained over 50% of the particles and appeared to lack a DARPin, so it was selected for an additional round of 3D classification without alignment. The resulting five classes (bottom row) show low resolution density in the GFP:DARPin region of the map.

**S1 Movie.**

Model of the DARPin-aldolase platform in complex with four spheres (radius=60 Å) anchored at the DARPin binding sites.

**S2 Movie.**

ResMap local resolution estimation of the DARPin-aldolase platform in complex with GFP.

## Reference

1. Henderson R. The potential and limitations of neutrons, electrons and X-rays for atomic resolution microscopy of unstained biological molecules. Q Rev Biophys. 1995;28(2):171-93.

2. Khoshouei M, Radjainia M, Baumeister W, Danev R. Cryo-EM structure of haemoglobin at 3.2 A determined with the Volta phase plate. Nat Commun. 2017;8:16099.

3. Brocchieri L, Karlin S. Protein length in eukaryotic and prokaryotic proteomes. Nucleic Acids Res. 2005;33(10):3390-400.

4. Kratz PA, Bottcher B, Nassal M. Native display of complete foreign protein domains on the surface of hepatitis B virus capsids. Proc Natl Acad Sci U S A. 1999;96(5):1915-20.

5. Coscia F, Estrozi LF, Hans F, Malet H, Noirclerc-Savoye M, Schoehn G, et al. Fusion to a homo-oligomeric scaffold allows cryo-EM analysis of a small protein. Sci Rep. 2016;6:30909.

6. Jensen GJ, Kornberg RD. Single-particle selection and alignment with heavy atom cluster-antibody conjugates. Proc Natl Acad Sci U S A. 1998;95(16):9262-7.

7. Wu S, Avila-Sakar A, Kim J, Booth DS, Greenberg CH, Rossi A, et al. Fabs enable single particle cryoEM studies of small proteins. Structure. 2012;20(4):582-92.

8. Liu Y, Gonen S, Gonen T, Yeates TO. Near-atomic cryo-EM imaging of a small protein displayed on a designed scaffolding system. Proc Natl Acad Sci U S A. 2018;115(13):3362-7.

9. Padilla JE, Colovos C, Yeates TO. Nanohedra: using symmetry to design self assembling protein cages, layers, crystals, and filaments. Proc Natl Acad Sci U S A. 2001;98(5):2217-21.

10. Jeong WH, Lee H, Song DH, Eom JH, Kim SC, Lee HS, et al. Connecting two proteins using a fusion alpha helix stabilized by a chemical cross linker. Nat Commun. 2016;7:11031.

11. Graille M, Stura EA, Corper AL, Sutton BJ, Taussig MJ, Charbonnier JB, et al. Crystal structure of a Staphylococcus aureus protein A domain complexed with the Fab fragment of a human IgM antibody: structural basis for recognition of B-cell receptors and superantigen activity. Proc Natl Acad Sci U S A. 2000;97(10):5399-404.

12. Brauchle M, Hansen S, Caussinus E, Lenard A, Ochoa-Espinosa A, Scholz O, et al. Protein interference applications in cellular and developmental biology using DARPins that recognize GFP and mCherry. Biol Open. 2014;3(12):1252-61.

13. Bartesaghi A, Merk A, Banerjee S, Matthies D, Wu X, Milne JL, et al. 2.2 A resolution cryo-EM structure of beta-galactosidase in complex with a cell-permeant inhibitor. Science. 2015;348(6239):1147-51.

14. Kudryashev M, Wang RY, Brackmann M, Scherer S, Maier T, Baker D, et al. Structure of the type VI secretion system contractile sheath. Cell. 2015;160(5):952-62.

15. Shoji S, Dambacher CM, Shajani Z, Williamson JR, Schultz PG. Systematic chromosomal deletion of bacterial ribosomal protein genes. J Mol Biol. 2011;413(4):751-61.

16. Yao Q, Lu Q, Wan X, Song F, Xu Y, Hu M, et al. A structural mechanism for bacterial autotransporter glycosylation by a dodecameric heptosyltransferase family. Elife. 2014;3.

17. Herzik MA, Jr., Wu M, Lander GC. Achieving better-than-3-A resolution by single-particle cryo-EM at 200 keV. Nat Methods. 2017;14(11):1075-8.

18. Ullmann A, Jacob F, Monod J. Characterization by in vitro complementation of a peptide corresponding to an operator-proximal segment of the beta-galactosidase structural gene of Escherichia coli. J Mol Biol. 1967;24(2):339-43.

19. Naydenova K, Russo CJ. Measuring the effects of particle orientation to improve the efficiency of electron cryomicroscopy. Nat Commun. 2017;8(1):629.

20. Hansen S, Stuber JC, Ernst P, Koch A, Bojar D, Batyuk A, et al. Design and applications of a clamp for Green Fluorescent Protein with picomolar affinity. Sci Rep. 2017;7(1):16292.

21. Zhou M, Li Y, Hu Q, Bai XC, Huang W, Yan C, et al. Atomic structure of the apoptosome: mechanism of cytochrome c- and dATP-mediated activation of Apaf-1. Genes Dev. 2015;29(22):2349-61.

22. Zhang Z, Liang WG, Bailey LJ, Tan YZ, Wei H, Wang A, et al. Ensemble cryoEM elucidates the mechanism of insulin capture and degradation by human insulin degrading enzyme. Elife. 2018;7.

23. Seeger MA, Zbinden R, Flutsch A, Gutte PG, Engeler S, Roschitzki-Voser H, et al. Design, construction, and characterization of a second-generation DARP in library with reduced hydrophobicity. Protein Sci. 2013;22(9):1239-57.

24. Yu H, Kohl A, Binz HK, Pluckthun A, Grutter MG, van Gunsteren WF. Molecular dynamics study of the stabilities of consensus designed ankyrin repeat proteins. Proteins. 2006;65(2):285-95

25. Kohl A, Binz HK, Forrer P, Stumpp MT, Pluckthun A, Grutter MG. Designed to be stable: crystal structure of a consensus ankyrin repeat protein. Proc Natl Acad Sci U S A. 2003;100(4):1700-5.

26. Amstutz P, Binz HK, Parizek P, Stumpp MT, Kohl A, Grutter MG, et al. Intracellular kinase inhibitors selected from combinatorial libraries of designed ankyrin repeat proteins. J Biol Chem. 2005;280(26):24715-22.

27. Kohl A, Amstutz P, Parizek P, Binz HK, Briand C, Capitani G, et al. Allosteric inhibition of aminoglycoside phosphotransferase by a designed ankyrin repeat protein. Structure. 2005;13(8):1131-41.

28. Kramer MA, Wetzel SK, Pluckthun A, Mittl PR, Grutter MG. Structural determinants for improved stability of designed ankyrin repeat proteins with a redesigned C-capping module. J Mol Biol. 2010;404(3):381-91.

29. Interlandi G, Wetzel SK, Settanni G, Pluckthun A, Caflisch A. Characterization and further stabilization of designed ankyrin repeat proteins by combining molecular dynamics simulations and experiments. J Mol Biol. 2008;375(3):837-54.

30. Pettersen EF, Goddard TD, Huang CC, Couch GS, Greenblatt DM, Meng EC, et al. UCSF Chimera--a visualization system for exploratory research and analysis. J Comput Chem. 2004;25(13):1605-12.

31. Mastronarde DN. Automated electron microscope tomography using robust prediction of specimen movements. J Struct Biol. 2005;152(1):36-51.

32. Kremer JR, Mastronarde DN, McIntosh JR. Computer visualization of three-dimensional image data using IMOD. J Struct Biol. 1996;116(1):71-6.

33. Zheng SQ, Palovcak E, Armache JP, Verba KA, Cheng Y, Agard DA. MotionCor2: anisotropic correction of beam-induced motion for improved cryo-electron microscopy. Nat Methods. 2017;14(4):331-2.

34. Grant T, Grigorieff N. Measuring the optimal exposure for single particle cryo-EM using a 2.6 A reconstruction of rotavirus VP6. Elife. 2015;4:e06980.

35. Kimanius D, Forsberg BO, Scheres SH, Lindahl E. Accelerated cryo-EM structure determination with parallelisation using GPUs in RELION-2. Elife. 2016;5.

36. Punjani A, Rubinstein JL, Fleet DJ, Brubaker MA. cryoSPARC: algorithms for rapid unsupervised cryo-EM structure determination. Nat Methods. 2017;14(3):290-6.

37. Emsley P, Cowtan K. Coot: model-building tools for molecular graphics. Acta Crystallogr D Biol Crystallogr. 2004;60(Pt 12 Pt 1):2126-32.

38. Kucukelbir A, Sigworth FJ, Tagare HD. Quantifying the local resolution of cryo-EM density maps. Nat Methods. 2014;11(1):63-5.

